# Four Decades of Breeding and Increasing Plant Density Management Reshaped Maize Root System Architecture

**DOI:** 10.64898/2025.12.10.693575

**Authors:** August C. Thies, Cintia Sciarresi, Slobodan Trifunovic, Douglas Eudy, Tony J. Vyn, Sotirios V. Archontoulis, Christopher N. Topp

## Abstract

Understanding how crop root system architecture (RSA) is influenced by genetics and management is vital for root-based improvement. We examined the impact of four decades of maize breeding on RSA using a panel of maize hybrids planted at different densities across six site-years in the Midwest. Root crowns were shovel-excavated and imaged via X-ray tomography, generating dozens of fine-grained 3D traits. Modern root systems were significantly smaller with fewer, thinner roots at higher density compared to older hybrids. However, the total amount of roots in the topsoil extrapolated across an acre was indistinguishable, revealing an equilibrium between crown size and density. Surprisingly, there was a 20% increase in the soil volume explored by individual modern hybrids, indicating increased overlap of root systems. We estimated modern hybrids share 43–46% of the topsoil with neighbors, compared to 14% in the oldest. Furthermore, when the oldest hybrids were grown at modern densities, their root crowns became more elliptical than modern hybrids, highlighting potential differences in avoidance response. Root systems of modern hybrids may be more intertwined and less competitive, revealing an adaptation to increased density stress in maize agriculture with implications for trait-based approaches to engineer more productive, efficient systems.

**HIGHLIGHT:** This study uses a new panel of maize hybrids to explore how decades of yield-focused breeding and advancements in planting density have indirectly impacted maize root systems.

## INTRODUCTION

Maize yield has steadily increased over the last century due to a combination of genetic improvement and management practices (Duvick 1977). One of the main drivers of historical maize yield gains has been the increase in plant density (Duvick *et al*. 2004). Concomitant to the adaptation to higher planting densities, other maize traits have also changed, such as increases in ear size and leaf angle (dos Santos *et al*., 2023; Elli *et al*., 2024; 2025), and decreases in anthesis-silking interval (Duvick *et al*. 2004; Wang *et al*. 2020; King *et al*., 2025). While many above-ground maize traits and their responses to yield selection have been researched, there have been fewer studies on below-ground maize traits. Continually rising planting densities make root responses to crowding a critical determinant of maize performance and management strategies.

Since maize breeding programs have prioritized grain yield and aboveground traits, any changes in root traits are presumed to have been a result of indirect selection. A prior growth simulation study estimated that increases in the steepness of root angles and deepness of root systems could increase water foraging and may have contributed to historical yield increases (Hammer *et al*. 2009). Yet, two subsequent empirical field studies have identified changes in root system architecture across a historical perspective with conflicting results on root system steepness (York *et al*. 2015; Ren *et al*. 2022), and whether any changes to root systems specifically influenced crop performance has been called into question (Messina *et al*. 2021). A more recent era study at V8 in large mesocosms found reductions in root system size and root-shoot-ratio in modern hybrids (Rinehart *et al*. 2024), but in a large field study that overlaps with this report, Sciarresi *et al*. (2024, 2025) found that the rate of growth in the topsoil and maximum depth has increased during breeding. Nonetheless, root traits remain one of the least explored dimensions of crop phenotypes, in part because field-ready methods to measure them are still evolving.

Due to the difficulty observing maize root system architecture complexity in the field, many prior studies have largely relied on combinations of simulations, controlled environments, or relatively coarse field methods such as measuring root pulling force or soil core sampling which can only provide plot-level data ( Wasson *et al*. 2014; Shao *et al*. 2021). Of late, image-based phenotyping methods have been developed that can estimate traits that would be more difficult, of poorer quality, or impossible to measure with conventional methods ( Grift *et al*. 2011; Das *et al*. 2015; Colombi *et al*. 2015; Seethepalli *et al*. 2021). The most advanced RSA phenotyping generates 3D models of field-excavated roots, including optical (Liu *et al*. 2021) and X-ray based methods (Jiang *et al*. 2019; Zeng *et al*. 2021; Duncan & Topp 2022). 3D root phenotyping has been shown to provide more statistical power to identify genotype-to-phenotype relationships by increasing heritability estimates and capturing more of the phenotypic space (Bray & Topp 2018; Shao *et al*. 2021; Hein *et al*. 2025).

To expand upon earlier studies of RSA and provide the most comprehensive evaluation of root crown phenotypes to date, we employed 3D root phenotyping on >1,700 field grown samples, collected over two years at three field locations in the Midwest United States. Using an advanced 3D root phenomics platform, TopoRoot+ (Ju *et al*. 2024), we digitally dissected the root crowns into constituent traits to provide a comprehensive evaluation of any changes to root system architecture that may have resulted from breeding for yield under increasing plant density. The germplasm represented over 30 years of plant breeding in Bayer’s legacy “era” hybrids, which underpinned yield gains of 38% during a time when average plant density nearly doubled, from 4.7 m^−2^ to 8.7 plants m^−2^ (Sciarresi *et al*. 2024). Best linear unbiased predictors (BLUPs) from across the six site-years provided little evidence of genetic gain for most root traits, including nodal root angles, which were unexpectedly (York *et al*. 2015; Ren *et al*. 2022) consistent across the eras. One remarkable exception was a steep increase (8%/year; 17% total) in convex hull volume, which measures the amount of soil explored by a root crown. This counterintuitive finding indicates that, rather than growing steeper and avoiding inter-row competition, the root systems of modern hybrids appear to be increasingly intertwined with their neighbors. Comparisons of era hybrid root systems at their historical or a common current density further supported existing evidence that hybrids adapted to lower densities have much smaller root systems when grown at higher density, but also that they change the 3D distribution of their root systems, which may influence contact with neighbors. Regardless of the mechanism, our work provides a comprehensive and granular view of the indirect effects of yield selection on maize root systems in a major breeding pool. We present evidence that the smaller individual root crowns in denser plantings are offset by the increased number of plants, resulting in a consistent amount of roots in the topsoil across the last four decades of maize agriculture, albeit in support of very different yield outcomes.

## MATERIALS AND METHODS

### Field experiment design

We designed a planting density experiment during 2021 and 2022 at three field sites across the US Corn Belt. The locations were Jerseyville, IL, West Lafayette, IN and Fraser, IA each year, for a total of six site-years. 11 hybrids from the Bayer Crop Science breeding program were chosen based on commercial relevance and germplasm representation from across four decades of breeding, ranging from 1983 to 2017 for the year of release (Supplementary Table 1). Up to three maize hybrids were selected from each of four eras (1980s ,1990s, 2000s and 2010s) and planted at the historical density of that hybrid 20,000, 25,000, 30,000 and 35,000 seeds/acre (corresponding to 4.7 plants m^−2^, 6 plants m^−2^, 7.3 plants m^−2^ and 8.7 plants m^−2^ respectively).

Each genotype was additionally planted at the higher, current density, 35,000 seeds/acre equivalent to 8.7 plants m^−2^. These field sites had randomized block design and three replications at each. They were rainfed, with conventional tillage, and the prior crop was soybean. Nitrogen was non-limited and ∼200 lbs/acre for each field. All field plots were 10 m wide with 8 rows and spacing at 0.76 m. Planting dates were staggered and for 2021 were April 15 (Jerseyville), May 2 (West Lafayette) and April 30 (Fraser). Planting dates for 2022 were May 10 (Jerseyville), May 11 (West Lafayette), and May 13 (Fraser). Precision planters were used and operated by Bayer Crop Science’s Field-Testing Organization. Tillage, fertilizing, weeding and other upkeep were managed by the local farmers, using standard farm practices per each region. Supplemental Figure 1 shows the location of the field sites where root crowns were collected, and root traits were observed.

### Root crown excavation

We shovel-excavated root crowns at growth stage R3, obtaining up to 3 samples from 2 reps for a total of 6 samples per era hybrid at each density treatment and field site. This totaled to ∼144 root crowns per field site for both the 2021 and 2022 field seasons for a total of 864 root crowns. The digs were staggered across three weeks to account for the travel time between sites and to better match the initial planting dates of each field site to obtain samples from plants at the same developmental stage. The excavated root crowns were then washed to remove excess soil and dried for 2 days at 28**°**C in a greenhouse. Root crown traits were extrapolated to a field-scale for certain comparisons and in those cases, we refer to the roots in the “topsoil” as those in the top 1-20 cm of soil due to the average depth they were collected at.

### Imaging and analysis

Washed root crowns were scanned via a Northstar X5000 X-Ray computed tomography system (North Star Imaging, MN, USA). Most of the root crowns were scanned for three minutes, with some taking longer depending on their size. 1800 radiographs at 10 fps were generated per crown, with a resolution of 109 µm. For further information see Duncan and Topp 2022. Scans were then exported as vertical (y-axis) image slices, thresholded using an automated algorithm, and converted to a skeletonized image and a point cloud image for trait analysis.

Further detail on these image analysis methods can be found in Shao *et al*. 2021. These 3D volumes were then analyzed using a custom computational script created by the Topp lab (https://github.com/Topp-Roots-Lab/3d-root-crown-analysis-pipeline; Shao 2021). This script produces over 100 different traits, such as total volume, total root length and root tip number. Additionally, a separate image analysis method called Toporoot was used on the root crown 3D volumes (https://github.com/Topp-Roots-Lab/TopoRoot; Zeng *et al*. 2021). This method creates a complete geometric skeleton of the root crowns, nearly free of errors. The skeleton is organized according to a topological hierarchy to distinguish between root types such as nodal roots and lateral roots and facilitates measures of each root in the root crown for distributions of measures across root types in a single root crown, including nodal root lengths, thicknesses, and emergence angles at each whorl. Examples of some of the root crown traits are shown in Figure 1.

**Figure 1:**
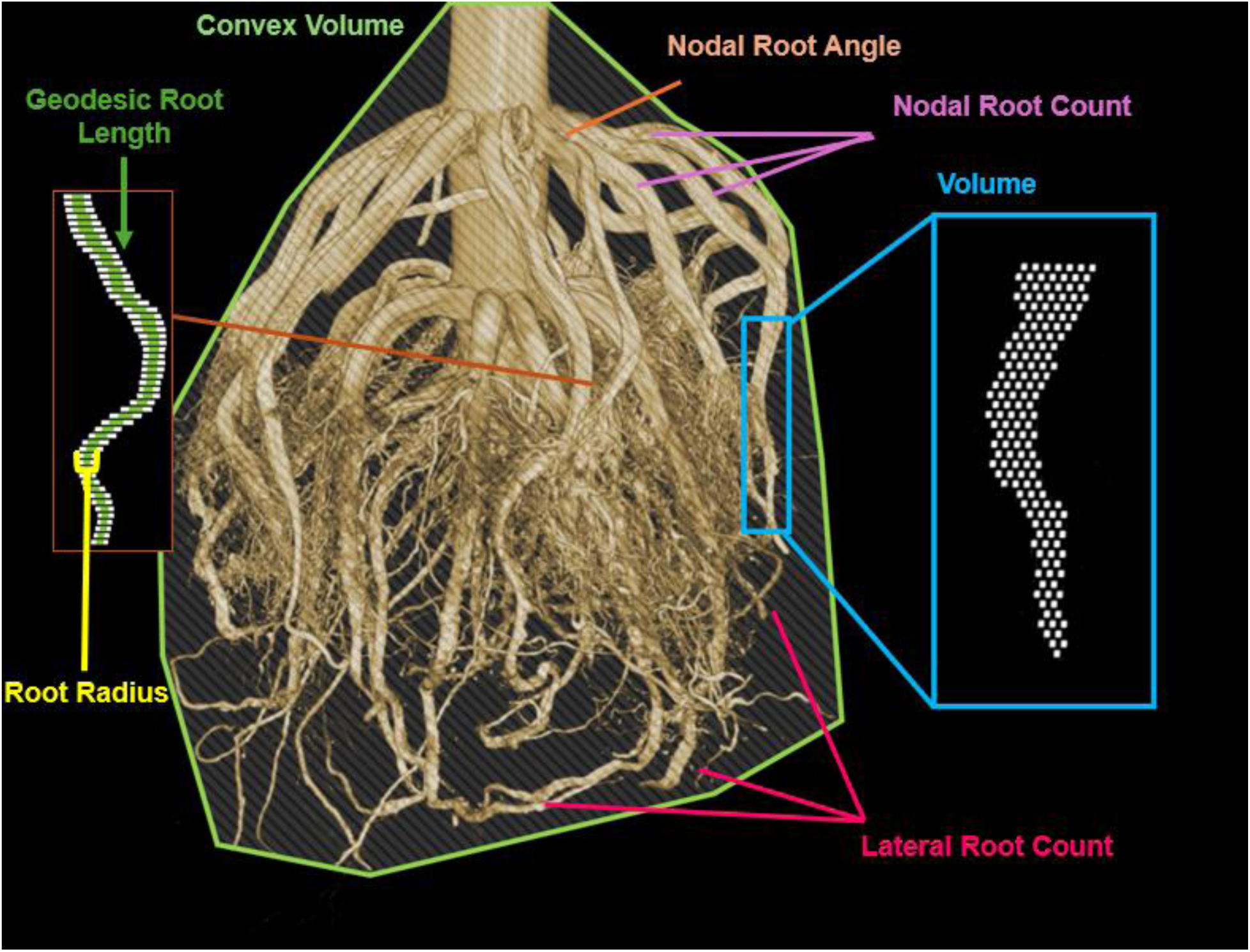
A reconstructed volume of a maize root crown with examples of root traits calculated from our custom computational pipeline and TopoRoot.

### Statistical analysis

The whole suite of root crown traits was then compared using statistical tests and multivariate analysis methods using R software version 4.5.0. For univariate analyses, outliers for traits were removed if they were outside three standard deviations from the average. ANOVA was performed on the trait once outliers were removed. Outliers were removed on a single-trait basis and were considered to be any trait values outside of 3 standard deviations for normally distributed traits and any values outside of 3*IQR for non-normal distributions. PCA was also performed on the genotypes, eras and locations in order to observe overall data trends. Additionally, linear mixed models were used to generate the best linear unbiased predictions (BLUPs) of different root crown traits.

### Genetic gain

The genetic gain across environments for the root traits and grain yield was estimated by fitting a linear mixed model, using the same model parameters described in Sciarresi *et al*. 2025: *“Year of release, the environment, and their interaction were considered fixed effects. The year of hybrid release was treated as a regression variable; the slope associated with year of hybrid release is the estimated genetic gain per year. Hybrid, hybrid within the environment, and block within the environment were considered random effects. In detail, the model was given as follows:*

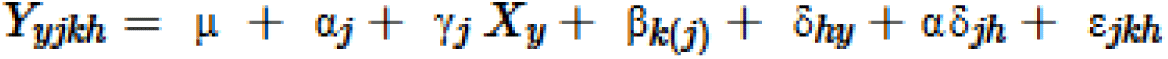

*where Y_yjkh_ is yield, RFV or maximum root depth for each year of hybrid release y, hybrid h, in a block k, nested within each environment j; μ is the common intercept; α_j_ is the fixed effect of the environment; ϒ_j_ is the slope (genetic gain) for year of hybrid release in environment j; β_k(j)_ is the random effect of block within the j environment; δ_hy_ is the random effect of the h hybrid within a year-of-release group, y; α_δjh_ is the random effect of the hybrid h within an environment j; and ε_jkh_ is the residual error*.

*Variance components for random effects were estimated by restricted maximum likelihood (REML). Coefficients for fixed effects were estimated by generalized least squares. The relative genetic gain (% per year) was calculated by dividing the absolute genetic gain of the trait by the mean genetic gain. Models were fit using the lmer function from the lme4 package from R software version 4.5.0 (Bates et al., 2015; R Core Team, 2022).”*

## RESULTS

### Yield and biomass increased in modern maize lines

Grain yield per environment averaged 14.4 Mg ha−1 (Fraser 2021), 13.7 Mg ha−1 (West Lafayette 2021), 13.6 Mg ha−1 (Jerseyville 2021), 13.9 Mg ha−1 (Fraser 2022), 12.9 Mg ha−1 (West Lafayette 2022) and 11.4 Mg ha−1 (Jerseyville 2022). Genetic gain for grain yield was positive across all locations and averaged 110 kg ha−1 year−1 (p < 0.001; Supplementary Table 2). Grain yield increased as planting density increased and for each subsequent era at all field sites (Supplementary Table 2). Aboveground biomass was 22.0 Mg ha−1 (Fraser 2021), 22.0 Mg ha−1 (West Lafayette 2021), 20.0 Mg ha−1 (Jerseyville 2021), 25.1 Mg ha−1 (Fraser 2022), 20.0 Mg ha−1 (West Lafayette 2022) and 19.8 Mg ha−1 (Jerseyville 2022). Average aboveground biomass increased as planting density increased and for each subsequent era at all field sites (Supplementary Table 3).

### Root crown volume was unchanged across eras

An advantage of our approach is that we can measure the root crown traits of individual plants as well as multiply by the per acre plant density to estimate the trait expression at field scale. We first focused on root crown size, which indicates relative energy investments of the plant into soil resource capture. Root crown volume is generated by summing all the 3D pixels, or voxels, in each root crown’s 3D volume and multiplying by the voxel size. Prior work established this estimate of root crown volume is highly correlated with physical biomass (∼90% across hundreds of samples) (Shao *et al*. 2021) and can be considered an accurate proxy. On average, increased plant density led to a 41% decrease in individual root crown volume, from an average of 870 cm^3^ at 4.7 plants m^−2^ for the 1980s era to 510 cm^3^ at 8.7 plants m^−2^ for the 2010s era (p < 0.0002; Figure 2a). However, when compared at the same density, there was little to no change in root crown volume across the era groups, suggesting that breeding was not acting on root crown size per se (p = 0.5159; Figure 2a). These trends are consistent across all six site-years (Supplemental Figure 2a) despite some environmental variability. Given that shoot biomass in our era study has experienced a genetic gain of 29.5% (Supplemental Table 3), we calculated the root-shoot-ratio by using the field-scaled root volume and comparing it to the aboveground shoot biomass (Supplemental Figure 6). Due to a constant per-plant root volume but increasing shoot biomass, there was a significant decrease of 17% in the root-to-shoot ratio across the eras due to breeding (p = .0166, Supplemental Figure 6), in agreement with prior work (Rinehart *et al*. 2024) and supporting a potential increased efficiency of root systems on a resource capture per energy investment scale (Messina 2021). There was no significant effect due to planting density (p = .6349, Supplemental Figure 6). We then asked what the per plot measurements scaled to across an acre for each era at either its historical density or current density. The total root volume in the top 1-20 cm (“topsoil”) of each field site for genotypes at their historical densities (p = .673; Figure 2) or at the current density (p = .5830, Figure 2) were not significantly different, indicating that the 1.75 fold increase in plant density from the mid-1980s to mid 2010s was balanced by smaller individual root crowns such that the amount of roots that have occurred in the topsoil during this time has been consistent at about 1.75 million cm^3^/acre.

**Figure 2:**
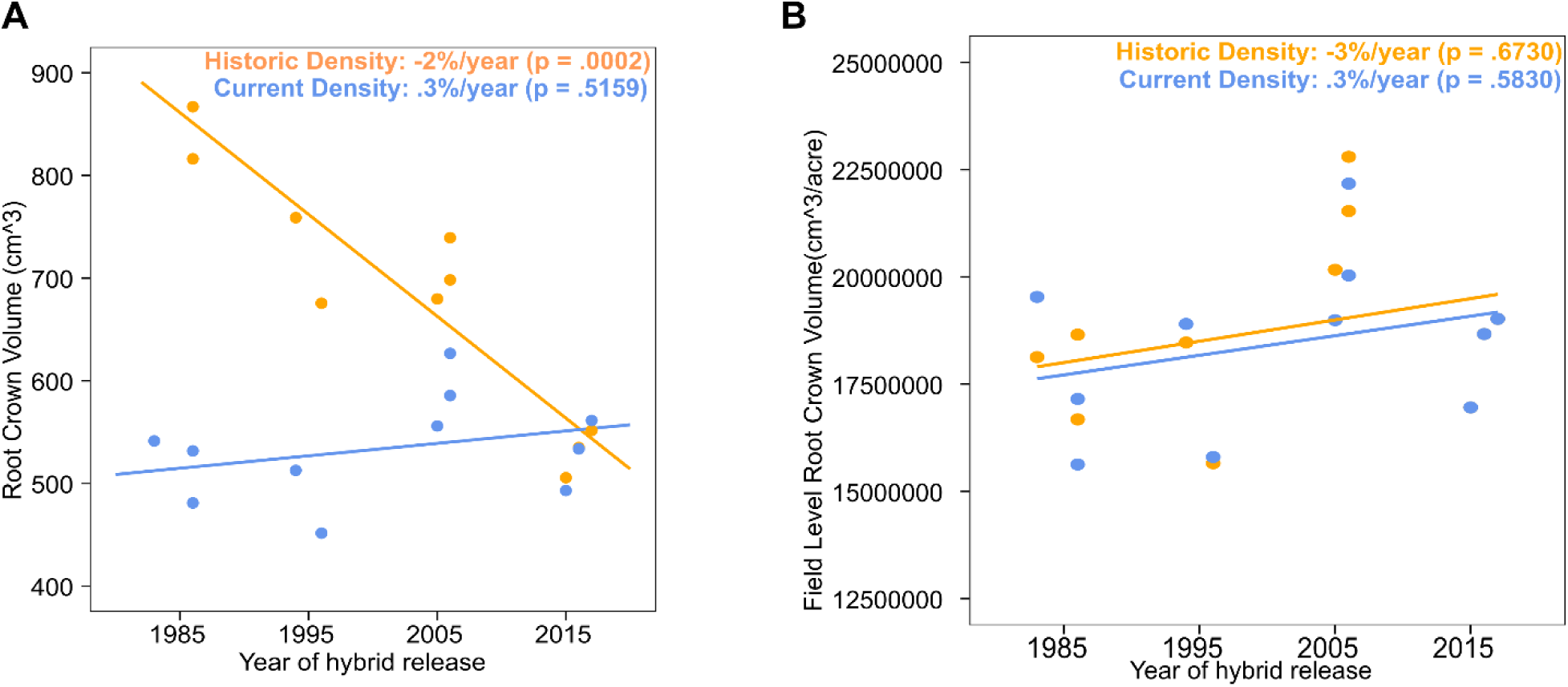
**A)** Best linear unbiased predictions (BLUPs) of maize root crown volume for the genotypes of different eras across all field sites in 2021 and 2022. The orange line is the historical density for a given era (ranging from 4.7 plants m-2 - 8.7 plants m-2) and shows a significant decrease as the planting density increases (−2.5%/year, **p = .0002**). The blue is for the current (8.7 plants m-2) planting density and is the genetic gain, showing no significant change in root crown volume (.3%/year, p = .5159). **B)** BLUPs for the root crown volume for genotypes on a field level scale when planted at their historic planting density, ranging from 20K/acre to 35K/acre. The estimated field level root crown volume when planted at historical density for a given era (ranging from 4.7 plants m^−2^ - 8.7 plants m^−2^) shows no significant change as the planting density increases (3%/year, p = .673). The estimated field level root crown volume at current (8.7 plants m^−2^) planting density shows no significant change in root crown volume each year (3%/year, p = .583).

### Root crown architecture has changed across eras

In addition to the total amount of roots, we queried how and where they were distributed, using a set of traits that collectively describe root system architecture. One key trait is the total number of root tips, which counts all the roots in the root system. The total number of root tips per crown decreased precipitously in more modern hybrids across historical plant densities, from an average of 4,300 to 3,100 root tips (29% reduction), but this difference was only marginally significant (p = .0576, Figure 3a). When grown at current density, root tip number was not significantly different across the eras (p = .2844, Figure 3a), suggesting this trait has been influenced more by planting density than genetics. Our system also measures the average root thickness (reported as radius) within individual root crowns, which showed significant decreases across historical plant densities, from 3.40 mm to 3.25 (4.5% reduction; p = .0124, Figure 3c).

**Figure 3:**
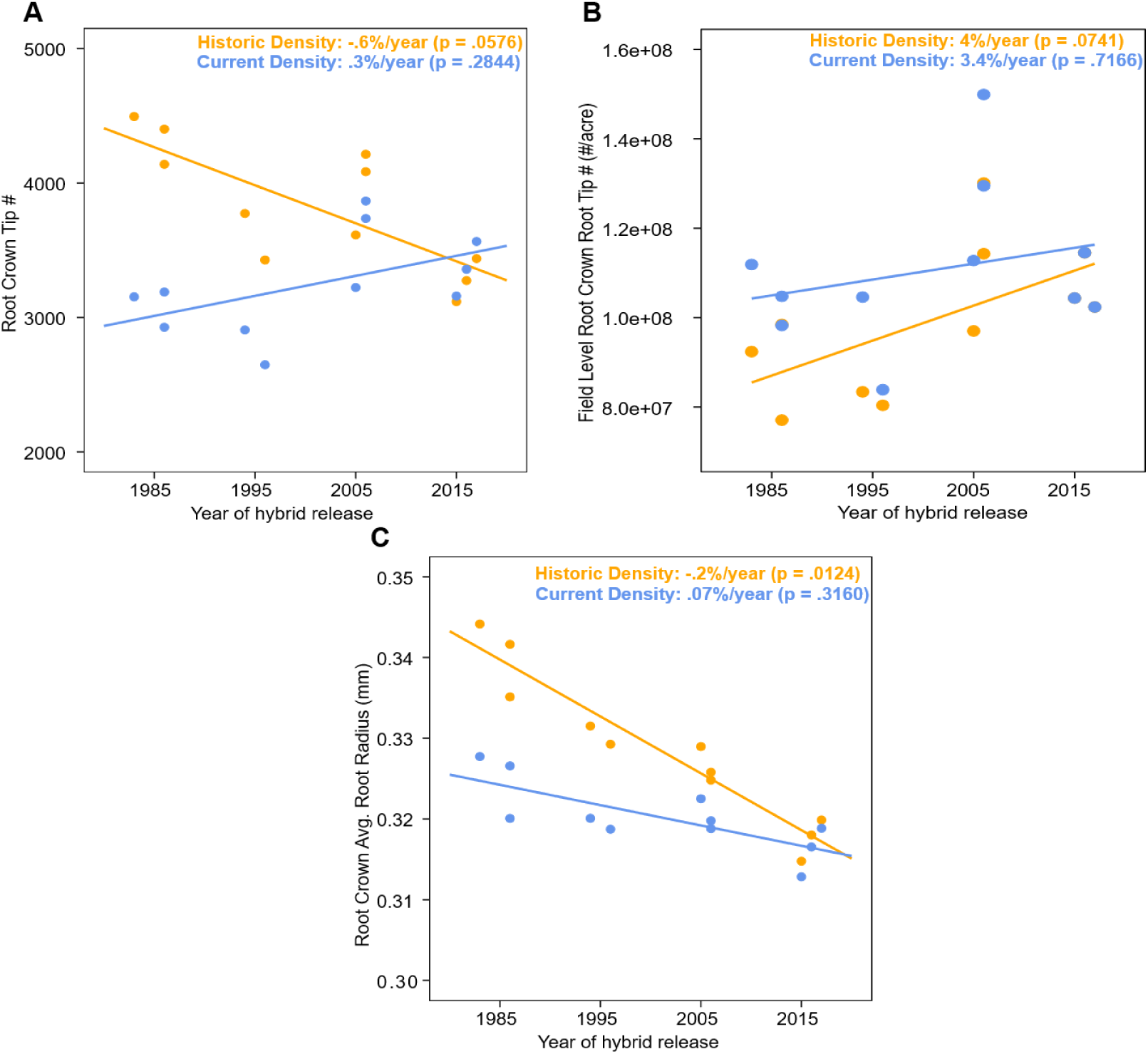
**A)** BLUPs of maize root crown tip # for the genotypes of different eras across all field sites in 2021 and 2022. The orange line is the historical density for a given era (ranging from 4.7 plants m^−2^ - 8.7 plants m^−2^) and shows a significant decrease as the planting density increases (−.6%/year, **p = .0576**). The blue is for the current (8.7 plants m^−2^) planting density and is the genetic gain, showing no significant change in root tip # across the eras (.3%/year, p = .2844). **B)** The BLUPs for root crown root tip number for genotypes on a field level scale when planted at their historic planting density, ranging from 20K/acre to 35K/acre. The plants at their historical density show a significant change as the planting density increases (4%/year, **p = .0741**). The estimated field level root crown root tip number for plants at current planting density shows no significant change in root tip number each year (3.4%/year, p = .7166). **C)** BLUPs of maize root crown average root radius for the genotypes of different eras across all of the field sites in 2021 and 2022. The plants at their historical density show a significant decrease as the planting density increases (−.2%/year, **p = .0124**). Plants at the current planting density show no significant change in average root crown radius (.07%/year, p = .3160).

Like root tip number, there were no significant differences in root thicknesses among the eras when grown at current density (p=.3160, Figure 3c). Despite per plant reductions in total roots per crown, there was an apparent large (20%) increase in the total number of roots in modern era hybrids when scaling across the field due to the greater number of plants, but it was only marginally significant (p=.0741, Figure 3b). This also seems to be a density effect as the average number of root tips among eras at field-scale was non-significant when eras were grown at current density (p = .7166, Figure 3b).

These results raised the question of how changes in the numbers and thicknesses of roots were apportioned into either nodal roots, which develop in circular whorls at the base of each leaf to form the major structure of the root system, or to lateral roots that emerge from nodal roots (first order) or other lateral roots (second order and beyond). To parse these root types in such dense and complex samples as excavated root crowns, we employed TopoRoot, a software which segments large and complex 3D root system models into their constituent parts by computing a nearly error-free geometric skeleton (TopoRoot, Zeng *et al*. 2021). A mild decrease in the total number of nodal roots per crown in more modern eras at their historical densities was statistically insignificant (p = 0.5871, Figure 4a), as when all eras were grown at current densities (p = 0.8082, Figure 4a) suggesting this trait was quite stable across the four decades of maize breeding in this germplasm pool. However, analysis of the stem thickness revealed a sharp reduction of 6.2% per year across this period (p = .0011, Supplemental Figure 3a), and when paired with the apparent lack of additional whorls of nodal roots (p = 0.5532), it reveals an increase in circumferential nodal root density (sometimes referred to as “occupancy”) of 8.4%. A 6.3% per year reduction (p = .0141, Figure 4d) in the average diameter of nodal roots in modern eras relative to older eras at historical densities comports with this observation, since nodal roots are typically at full occupancy and constrained to one plane in each whorl. Differences in nodal root thickness was indistinguishable when eras were grown at the same, current, density (p = .9238, Figure 4d). Yet the average length of each nodal root in the crown showed a striking genetic gain of 2.6% per year (11% total increase) in modern eras (p = .0283, Figure 4c) supporting indirect selection for this trait. Overall, there appears to have been allometric shifts in nodal rooting to become relatively more dense, thinner, but longer in modern germplasm.

**Figure 4:**
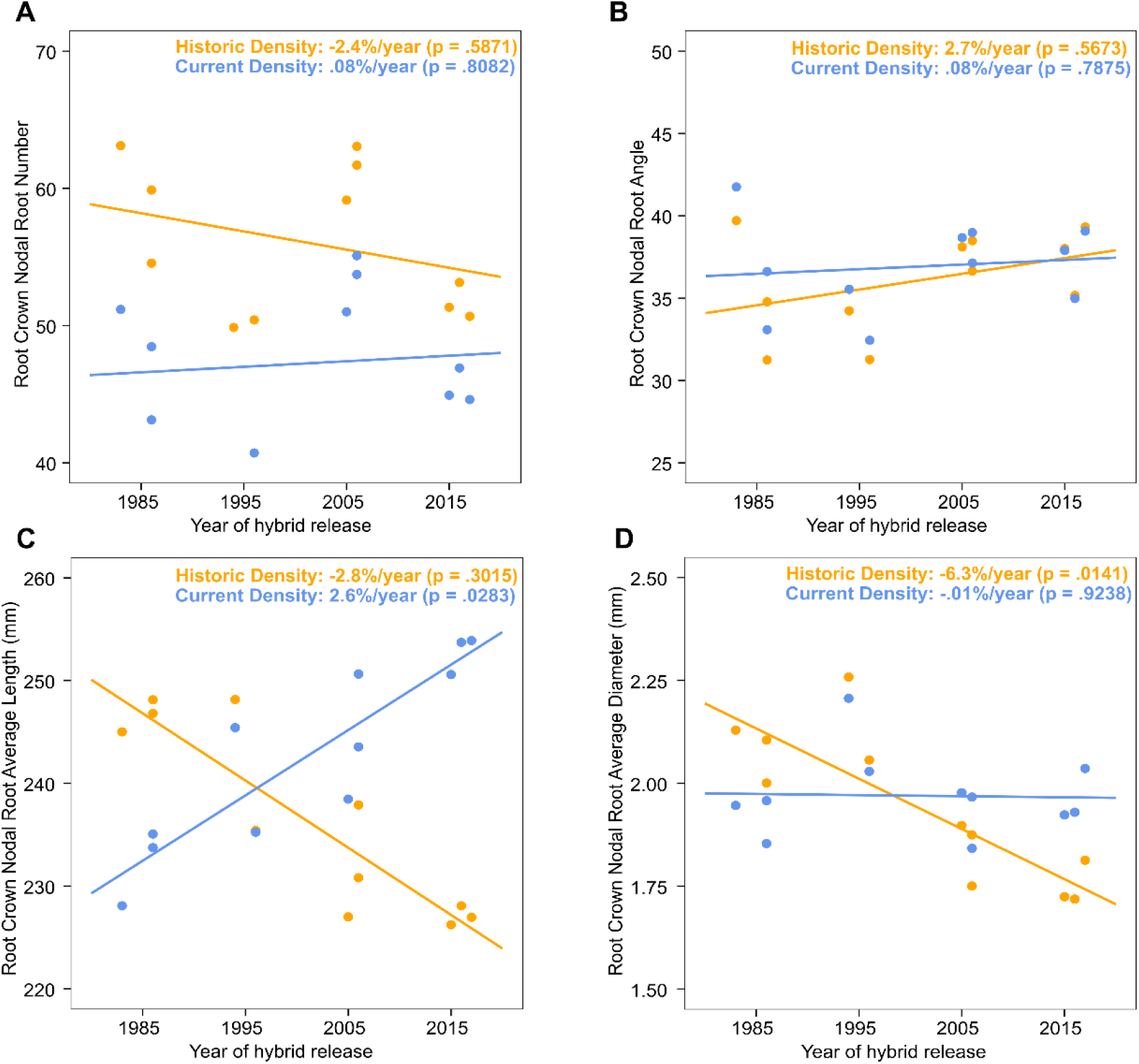
**A)** BLUPs of maize root crown nodal root # for the genotypes of different eras across all field sites in 2021 and 2022. The orange line is the historical density for a given era (ranging from 4.7 plants m^−2^ - 8.7 plants m^−2^) and shows no significant change as the planting density increases (−2.4%/year, p = .5871). The blue is for the current (8.7 plants m^−2^) planting density and is the genetic gain, showing no significant change in root tip # across the eras (.08%/year, p = .8082)**. B)** BLUPs of maize nodal root angle for the genotypes of different eras across all of the field sites in 2021 and 2022. The plants at their historical density show no significant change as the planting density increases (2.7%/year, p = .5637). The plants at current planting density show no significant changes in nodal root angle across the eras (.08%/year, p = .7875). **C)** BLUPs of maize nodal root length for the genotypes of different eras across all of the field sites in 2021 and 2022. The plants at their historical density show no significant decrease as the planting density increases (−2.8%/year, p = .3015). Plants at the current planting density show a significant change in nodal root length across the eras (2.6%/year, **p = .0283**). **D)** BLUPs of maize root crown nodal root thickness for the genotypes of different eras across all of the field sites in 2021 and 2022. The plants at their historical density show a significant decrease in nodal root diameter as the planting density increases (−6.3%/year, **p = .0141**). The plants at current planting density show no significant change in nodal root diameter over the eras (−.01%/year, p = .9238).

Examination of lateral root number revealed a large and significant decrease of 50% (p = .0028, Figure 5a) in modern eras, consistent with the overall reduction in root numbers (Figure 3a). There was a marginally significant (p = 0.0883, Figure 5b) genetic gain for lateral root length, but this was insignificant across eras at their historical densities, suggesting the gain simply maintained average lateral root length as density increased.

**Figure 5:**
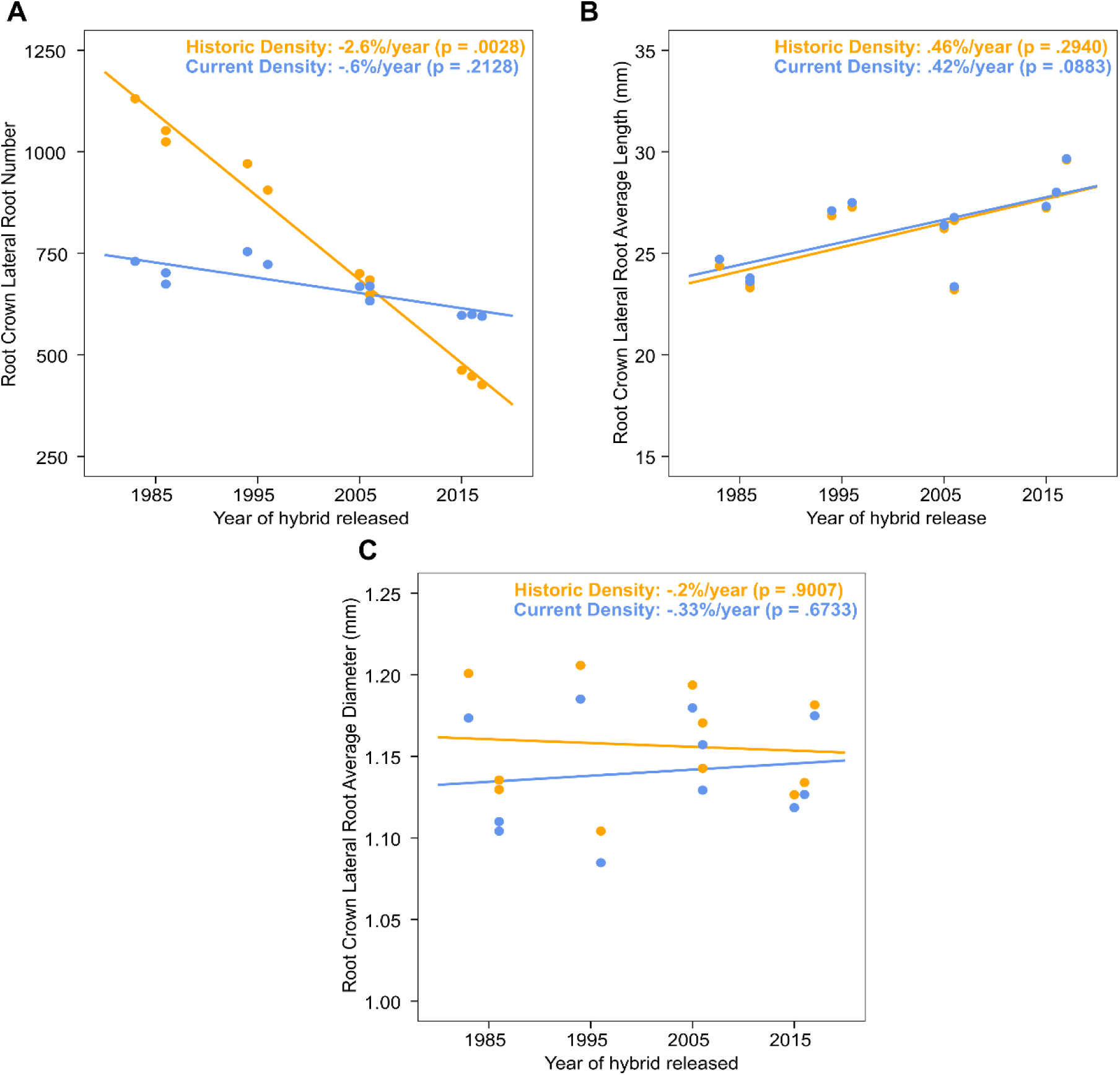
**A)** BLUPs of maize lateral root number for the genotypes of different eras across all field sites in 2021 and 2022. The orange line is the historical density for a given era (ranging from 4.7 plants m-2 - 8.7 plants m-2) and shows a significant decrease as the planting density increases (−2.6%/year, **p = .0028**). The blue is for the current (8.7 plants m-2) planting density and is the genetic gain, showing no significant change in lateral root count across the eras (−.6%/year, p = .2128). **B)** BLUPs of average maize lateral root length for the genotypes of different eras across all field sites in 2021 and 2022. The plants at their historical density show no significant change as the planting density increases (.46%/year, p = .2940). The plants at the current planting density show no significant change in average lateral root length over the eras (.42%/year, p = .0883). **C)** BLUPs of average maize lateral root thickness for the genotypes of different eras across all field sites in 2021 and 2022. The plants at their historical density show no significant decrease as the planting density increases (−.2%/year, p = .9007). The plants at the current density show no significant change in lateral root diameter across the eras (−.33%/year, p = .6733).

### Root crown geometry is different in modern maize, increasing root zone overlap of neighboring plants

Nodal root angle has been widely reported to be stress responsive and has also been observed to have been indirectly selected for during maize breeding (Ren *et al*. 2022 & York *et al*. 2015). Unlike prior works that used 2D analysis to assess overall root crown angle, our TopoRoot pipeline estimated the emergence angle of every identified nodal root in the root crown. Strikingly, in our experiment, the average of all nodal root angles did not vary significantly due to either breeding (p = .5673, Figure 4b) or as planting density increased (p = .7875, Figure 4b).

Another key trait is convex hull volume, which estimates the total space that a root crown encompasses in the soil. Despite an essentially consistent nodal root emergence angle of all eras at any density, BLUPs showed a large (30%) and significant increase in root crown convex volume in more modern maize varieties (from 10,000 cm^3^ to 13,000 cm^3^; p = 0.0373; Figure 6). Like root crown volume, convex volume had a 25% decrease as planting density increased (p = .0020; Figure 6) from 20,000 cm^3^ to 15,000 cm^3^. This suggests that despite smaller volume, more modern root crowns were exploring more soil space on a per-plant-basis when compared to earlier lines. When measuring on a field scale, root crown convex volume was significantly increased due to greater planting density in more modern maize varieties by 43%, from 3.5e+08 cm^3^/acre to 5.0e+08 cm^3^/acre (p = .0439; Supplemental Figure 5). Despite the similar amounts of root crown volume, progressively modern root crowns explored more of the upper 1-20 cm of soil space in each field. As an estimate of thoroughness of this exploration, we calculated solidity by dividing root volume by the convex volume, providing the denseness of the root distribution in the crown. On a field level, solidity was shown to decrease as planting density increased (p = .0509, Supplemental Figure 4) and did not have any significant change through the eras (p = .3068, Supplemental Figure 4). These observations were counterintuitive to the prevailing notion that maize root systems generally maintained their individual space relative to neighbors to avoid a competition response (Hammer 2009, Messina 2021, Ren 2022) but instead suggested they have been increasingly overlapping their root zones.

**Figure 6:**
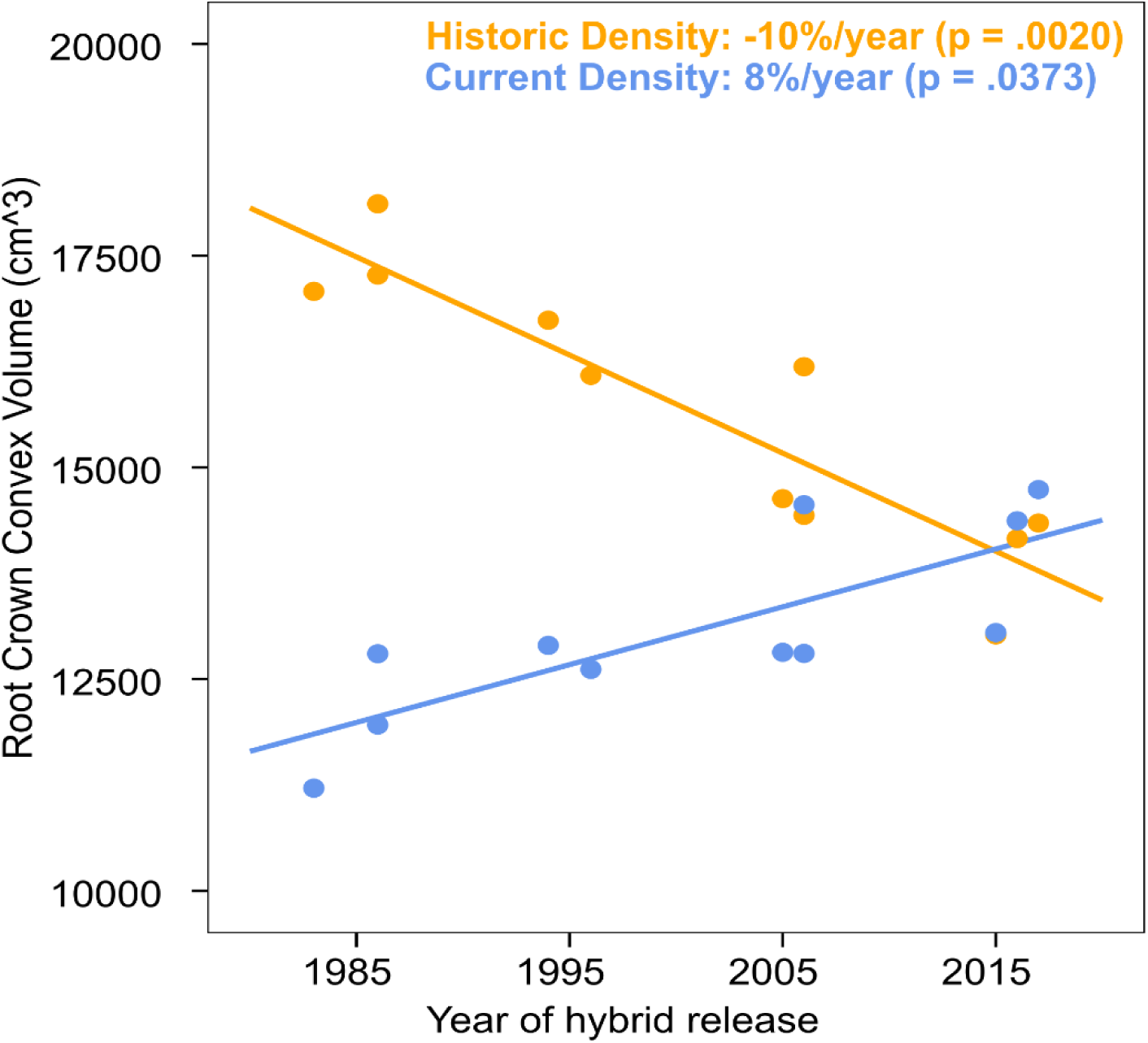
**A)** BLUPs of maize root crown convex volume for the genotypes of different eras across all field sites in 2021 and 2022. The orange line is the historical density for a given era (ranging from 4.7 plants m-2 - 8.7 plants m-2) and shows a significant decrease as the planting density increases (−10%/year, **p = .0020).** The blue is for the current (8.7 plants m-2) planting density and is the genetic gain, showing a significant increase in root crown convex volume across the eras (8%/year, **p = .0373**).

To explore this idea, we sought to estimate the geometry of individual root crowns relative to the average space between neighbors at a given plant density by evaluating their cylindricity around the axis of gravity (z), which is calculated from a principal component analysis (PCA) on (x, y) of the 3D point cloud and the ratio between PC2 variance and PC1 variance, referred to as “Root Crown Football” in our pipeline. This trait thus varies from 1 to 0, with 1 reporting a completely cylindrical root crown and 0 reporting an essentially “flat” 2-dimensional shape on either the x or y axis. The mean cylindricity of 1980 era root crowns planted at the current density (8.7 plants m^−2^) was lower than both the 1980 era planted at their historic density (4.7 plants m^−2^) and the 2010 era at 8.7 plants m^−2^, indicating the increase in plant density of non-density adapted hybrids resulted in a flattening of root crown geometry. To compare the distributions of root crown cylindricity, a two-sample Kolmogorov-Smirnov (KS) test was performed. The distribution mean of cylindricity for 1980s/4.7 plants m^−2^ was significantly lower than the distribution mean for the 1980s/8.7 plants m^−2^ (KS test, p = 0.0428) and lower than the mean for 2010s/8.7 plants m^−2^ (two-sample KS test, p = 0.0828). The distribution mean of cylindricity for 8.7 plants m^−2^ and 1980s/4.7 plants m^−2^ were not significantly different (KS test, p = 0.8497). Figure 7 shows the distributions of the root crown cylindricity trait among the earliest and most modern root crowns at their various planting densities. The findings that high planting density reduces the cylindricity of root crowns in older germplasm, but that modern root crowns maintain their relative cylindricity in an identical density, suggest a potential dampening of a root neighbor-avoidance response during maize breeding.

**Figure 7:**
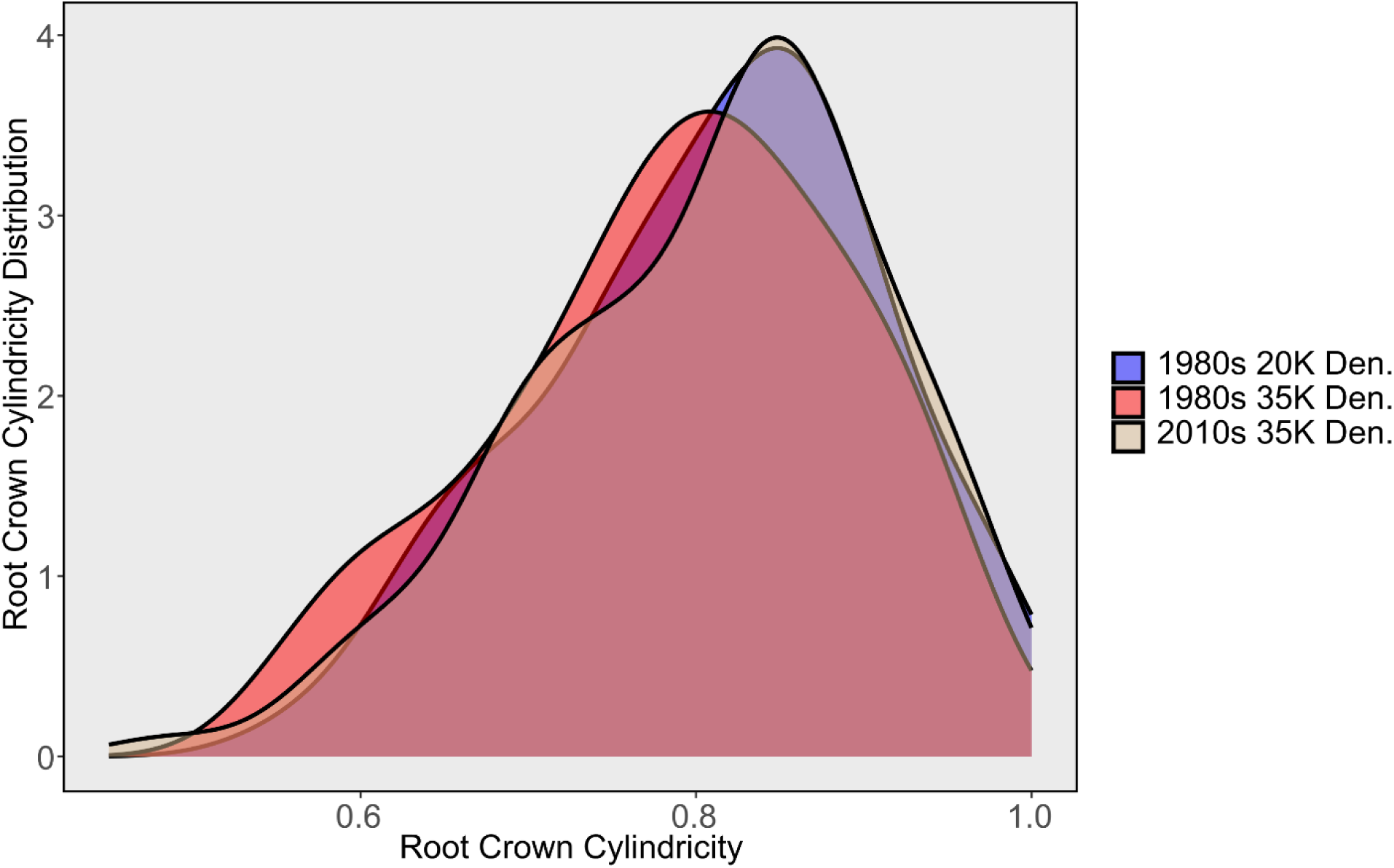
Distribution of the cylindricity trait of root crowns for the 1980s root crowns at both densities and the 2010s at 8.7 plants m^−2^ density. Values closer to 1 are more cylindrical whereas values closer to 0 are more flattened on one axis. The distribution mean of cylindricity for 1980s/8.7 plants m^−2^ is significantly lower than the distribution mean for the 1980s/4.7 plants m^−2^ (two-sample Kolmogorov-Smirnov (KS) test, **p = 0.0428**), and also lower than the distribution mean for 2010s/8.7 plants m^−2^ (two-sample KS test, p = 0.0828). The distribution means of cylindricity for 2010s/8.7 plants m^−2^ and 1980s/4.7 plants m^−2^ were not significantly different (two-sample KS test, p = 0.8497).

We were next interested in how these changes may affect the physical overlap of root systems across the eras. An estimate of the width of a given root crown can be found by calculating the maximum root system diameter among all horizontal slices of the model. Figure 8 shows the distributions of maximum width for the earliest and most modern root crowns at their various planting densities. The distribution mean of max horizontal diameter for 1980s/4.7 plants m^−2^ was significantly greater than the mean for the 1980s/8.7 plants m^−2^ (two-sample KS test, p < 0.001) and higher than the mean for 2010s/8.7 plants m^−2^ (two-sample KS test, p < 0.001). The distribution mean of max horizontal diameter for 8.7 plants m^−2^ and 1980s/8.7 plants m^−2^ was also significantly different (two-sample KS test, p = 0.0216).

**Figure 8:**
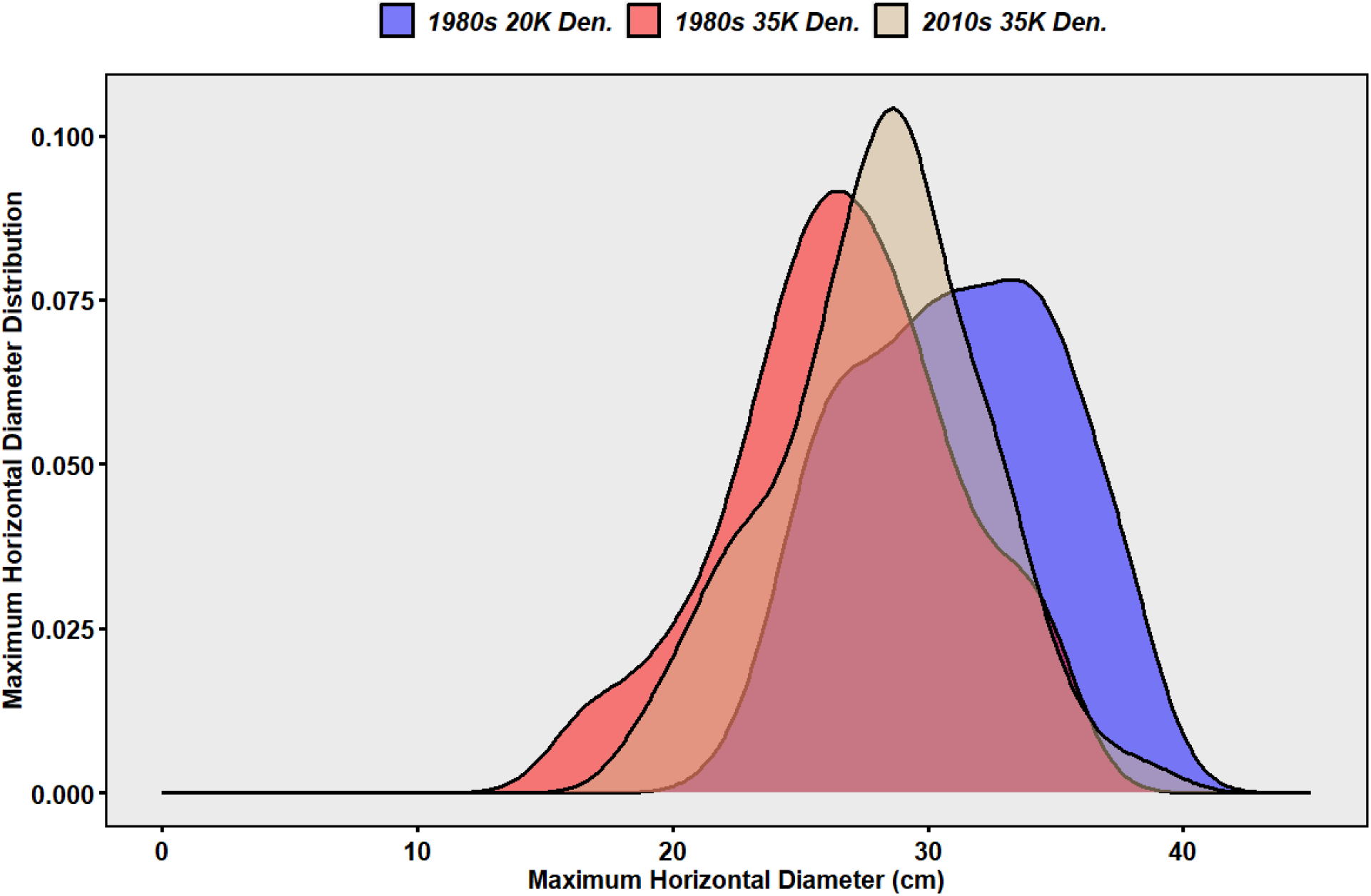
Distribution of the maximum horizontal diameter trait of root crowns for the 1980s root crowns at both densities and the 2010s at 8.7 plants m^−2^ density. The distribution mean of max horizontal diameter for 1980s/4.7 plants m^−2^ was significantly greater than the mean for the 1980s/8.7 plants m^−2^ (two-sample Kolmogorov-Smirnov (KS) test, p < 0.001) and also higher than the mean for 2010s/8.7 plants m^−2^ (two-sample KS test, **p < 0.001**). The distribution mean of max horizontal diameter for 2010s/8.7 plants m^−2^ and 1980s/8.7 plants m^−2^ was also significantly different (two-sample KS test, p = 0.0216).

The average maximum width for the root crowns from the 1980s era at 4.7 plants m^−2^ is 31.12 centimeters, for the 1980s era at 8.7 plants m^−2^ is 26.70 centimeters and for the 2010s era at 8.7 plants m^−2^ is 28.07 centimeters. The average row spacing between plants for 4.7 plants m^−2^ is 27 cm and for 8.7 plants m^−2^ is 15 cm. An estimate of overlap between root crowns at each planting density was calculated by subtracting the average row spacing between plants from the max width and multiplying by the cylindricity. This resulted in an average maximum root crown overlap of 4.4 cm, 11.5 cm and 12.8 cm for the 1980s/4.7 plants m^−2^, 1980s/8.7 plants m^−2^ and 2010s/8.7 plants m^−2^ respectively. Comparing this as a percentage of the total width resulted in ranges of 12.5-14%, 33-37% and 43-46% for 1980s/4.7 plants m^−2^, 1980s/8.7 plants m^−2^ and 2010s/8.7 plants m^−2^ root crowns respectively. Figure 9 provides an illustration of this root crown overlap for the 1980s/4.7 plants m^−2^ and 2010s/8.7 plants m^−2^ root crowns. Since we did not record the direction of the root crowns in the field, the lower and upper ends of the estimate account for the root crowns being more flattened perpendicular or parallel to the row respectively. Increased planting density was shown to decrease maximum root crown width in older germplasm, but modern root crowns are relatively wider, and thus likely had a greater proportion of their root crown overlapping neighboring root zones, indicating a possible mechanism of adaptation to high density that merits further study.

**Figure 9:**
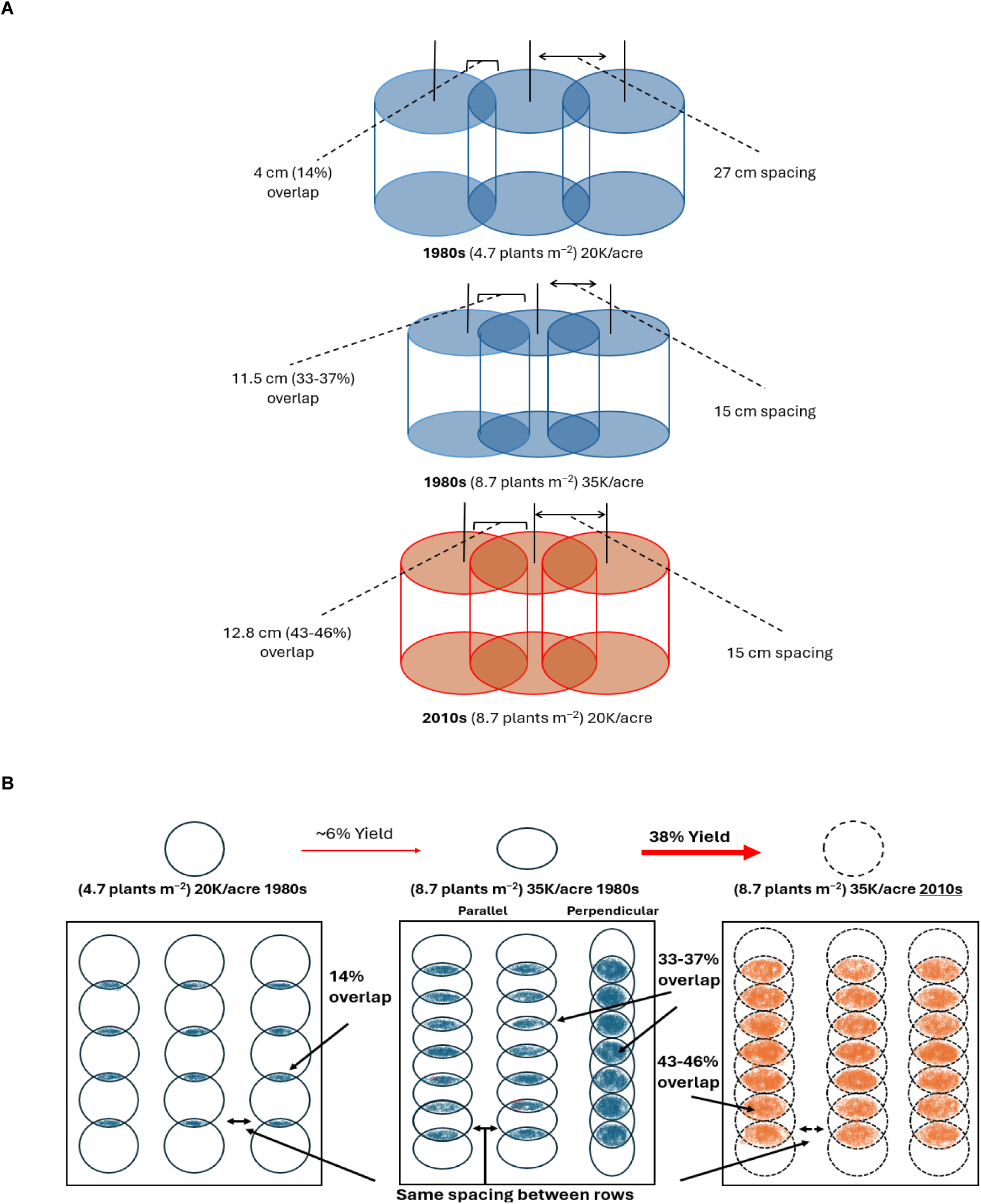
**A)** Illustration of how root crowns in the field are spaced out and intercalate at various densities from a profile view. **B)** Illustration of how root crowns in the field are spaced out and intercalate at various densities from an above view, with the 35K/1980s showing an additional parallel vs. perpendicular view to demonstrate the possible range of intercalation.

## DISCUSSION

### The root systems of modern hybrids have adapted to share more space in the soil than their historical counterparts

Greater planting densities led to a decrease in both root crown volume and root crown convex volume of individual plants (Figures 2 & 6), consistent with expectations under increased belowground competition (Jiang *et al*. 2013, Yu *et al*. 2019 & Shao *et al*. 2024). When examining breeding effects at the high density, root crown volume itself did not differ across eras (Figure 2). However, when all eras were planted at the same high density of 8.7 plants m⁻², the convex volume of individual crowns increased in more modern hybrids while root crown volume remained unchanged (Figures 2 and 6). This indicates that modern hybrids have a greater capacity to coexist with neighboring roots in the soil. Given that planting density has steadily increased over the last century (Duvick & Cooper, 2004), this increase in convex volume may reflect an architectural adaptation to crowding.

In line with this interpretation, yield and aboveground biomass per acre increased in later eras and at higher planting densities across all field sites (Supplementary Tables 1 & 2). Thus, the root systems of modern hybrids continue to support greater aboveground productivity despite more crowded conditions. When considered at a field scale, total root crown volume did not differ among eras or planting densities (Figure 2b), consistent with a decrease in root-to-shoot ratio. Likewise, total root number in the top 0–20 cm remained stable across eras when planting density was controlled (Figure 3b). Other work from the same Bayer era project (Sciarresi *et al*. 2024; Sciarresi *et al*. 2025) reported increases in both deep root biomass and the rate of root system expansion below the topsoil. These findings offer a mechanistic explanation for how the efficiency of the root system may have increased even though the amount of root volume in the topsoil did not change in our study. Strikingly, total root crown convex volume at the field scale increased across more modern lines (Supplemental Figure 5), and given the lack of change in root volume, this points to an architectural shift toward greater overlap of root systems under modern planting densities.

### Changes in root classes from planting density stress

Other components of RSA may also contribute to the observed improvements in aboveground productivity. Average thickness and number of roots in a crown decreased under higher planting densities (Figure 3), consistent with previous work (York *et al*. 2015; Shao *et al*. 2024). Both stem thickness and the number of whorls above the soil surface were reduced at higher density (Supplemental Figure 3). The proportional changes suggest these factors could explain the reduction in nodal root number, as stem thickness decreased by 16% (p = 0.011).

Nodal root length did not respond significantly to planting density (Figure 4c), while nodal root thickness decreased (Figure 4d). With nodal root thickness remaining constant and stem diameter reduced, fewer nodal roots could form per whorl; and because whorl number also did not increase, total nodal root number declined. These observations suggest that reductions in stem thickness are the primary driver of declines in nodal root number. Despite this, circumferential nodal root density increased by 8.4%, because the reduction in nodal number was proportionally smaller than the reduction in stem thickness.

In contrast to nodal roots, lateral root number was not affected by planting density (Figure 5a), and lateral root length and thickness were also largely unchanged, except for a slight reduction in thickness in the 2000s era (Figures 5a & 5c). York *et al*. (2015) reported reduced lateral root length under high density, which may reflect methodological differences, for example sampling a subset of lateral roots in their study versus measuring entire crowns here, or genetic differences in the germplasm studied. Nodal root angle also did not respond significantly to planting density (Figure 4b), which differs from previous reports showing changes in both directions. York *et al*. (2015) found shallower angles in earlier lines and steeper angles in more modern ones, whereas two other studies observed a reduction in root system width in more modern lines (Ren et al. 2022, Messina et al. 2021).

### Changes in root classes from breeding

Nodal root number, stem thickness, whorl number, and nodal root angle did not differ across eras (Figures 4 & S3), in contrast to some expectations based on earlier studies. York et al. 2015 showed that more modern root crowns had steeper nodal roots, but the lines they compared included much earlier eras. Furthermore, while Ren et al. 2022 studied germplasm more similar in age to ours, they focused on inbreds from a Chinese germplasm pool with unknown relation to our material. Nevertheless, in our study, modern hybrids exhibited significantly longer nodal roots while maintaining similar thickness (Figures 4c & 4d). Lateral roots also showed a small decrease in number and an increase in length in more modern lines (Figures 5a & 5b), similar to patterns reported in York et al. 2015.

The “steep, cheap, and deep” (SCD) ideotype (Lynch 2013) is relevant in the context of planting density stress. Although steepness did not change with density in our study, density-induced reductions in root volume, nodal roots, and stem thickness resemble a “cheaper” root response. Supporting this possibility, soil core and root front velocity data from Sciarresi et al. (2024, 2025) found that roots grew deeper and faster in more modern lines and at higher planting densities, which led to increased root mass at depth. If high planting density elicits responses similar to nutrient stress, RSA changes could reflect a coordinated response to increased resource competition. When considering breeding effects alone, we observed no dominant changes in steepness or volume across eras; however, more modern maize had longer nodal and lateral roots as well as greater convex volume. These patterns may indicate a shift toward deeper rooting as an adaptation to increased planting density, although the absence of low-density plantings for the most modern lines limits our ability to disentangle breeding effects from density responses. Overall, the data suggest that modern maize employs different strategies than earlier hybrids to cope with density stress.

### RSA Shape, Breeding Effects, and Potential Root–Root Interactions

The increased convex volume in modern hybrids reflects clear changes in root crown shape. Our 3D phenotyping identified breeding trends in both cylindricity and maximum crown width that describe how this shape is distributed in the soil. The 1980s era hybrids were nearly cylindrical around the depth axis at their historical density of (4.7 plants m⁻²), but became much more flattened when grown at modern, high density (8.7 plants m⁻²), potentially in response to crowding by neighbors. In contrast, the 2010 era hybrids were nearly cylindrical under the same density (Figure 7). Similarly, maximum crown width was greater in the 2010s era than in the 1980s era at 8.7 plants m⁻², indicating that modern crowns extend further laterally even when overall volume is unchanged (Figure 8). Combined with longer nodal and lateral roots, these patterns show that modern crowns explore a larger soil domain despite the constraints of higher planting density.

These architectural shifts have implications for how root systems interact belowground. Historically, the root crowns of hybrids grown in the 1980s had an average overlap of only 14%, but with increased density management this value climbed to 43-46% in modern hybrids (Figure 9). Whereas earlier-era crowns changed their root distributions to become more flattened under high density, the near cylindricity of modern hybrids at high density may indicate reduced aversion to neighboring roots and a greater propensity for intercalation compared to earlier lines. Because crown orientation was not recorded at excavation, we cannot determine whether flattening represented directional avoidance or possibly even alignment with the planting row.

However, even under extreme geometric assumptions of parallel flattening to maximize overlap or perpendicular to minimize it, modern hybrids consistently showed greater potential overlap with neighbors (43–46%) than earlier hybrids (34–37%; Figure 9) at high density, due to their larger convex hull areas. Given the conical structure of maize crowns, such root zone overlap likely increases with depth, emphasizing the extent to which modern germplasm shares crowded soil spaces.

### Future directions

To better understand the extent to which our observations describe broadly common effects of indirect selection on root systems during maize breeding, additional era panels representing other historically important germplasm could be evaluated across environments. Genetic mapping would link architectural traits to specific loci, and with the addition of haplotype and functional analysis, could pinpoint genetic variants that influence these traits. Additionally, testing modern lines at lower planting densities would reveal which RSA traits represent fixed adaptation or plasticity to increased density (Bernhard & Below 2020), perhaps by altering stress perception or response. Finally, in future studies, marking crown orientation during excavation could clarify whether differences in cylindricity and flattening response indicate neighbor avoidance, attraction, or some other environmental factor.

## Conclusion

This study provides a detailed view of how maize RSA responds to planting density and how breeding programs have shaped these responses. Higher density reduced root volume, convex volume, maximum crown width, cylindricity, and nodal root number. More modern lines exhibited greater soil exploration while maintaining the same volume, likely due to longer nodal and lateral roots and retention of crown shape under crowding. Modern crowns appear to intertwine more extensively with neighbors, indicating an improved response to density stress. Continued study of these traits will be important for breeding maize that sustains yield under rising global demand and increasingly crowded production systems.

## SUPPLEMENTARY DATA

Supplemental Figure 1: Map of all field sites and experiment types during the 2020-2022 field seasons. Sites where root traits were collected are marked with a red circle.

Supplemental Figure 2: Best linear unbiased estimates (BLUEs) for maize root crown volume & convex volume for the genotypes of different eras at each individual field site in 2021 and 2022.

Supplemental Figure 3: BLUPs of maize stem thickness for the genotypes of different eras across all field sites in 2021 and 2022.

Supplemental Figure 4: BLUPs of maize root crown solidity for the genotypes of different eras across all field sites in 2021 and 2022.

Supplemental Figure 5: The root crown convex volume for genotypes on a field level scale when planted at their historic planting density,

Supplemental Figure 6: Split violin plot of the root-shoot ratio of current planting density versus historical density.

Supplementary Table 1: Maize hybrids for each year of release are grouped in their different eras.

Supplementary Table 2: Average yields of each era at each field site for 2021 & 2022.

Supplementary Table 3: Average aboveground biomass of each era at each field site for 2021 & 2022.

## AUTHOR CONTRIBUTIONS

AT, and CNT: conceptualization; AT, CS, SVA, and CNT: methodology; AT: formal analysis; AT: software; AT, CS, SVA, and CNT: investigation; SVA, DE, TJV, ST, and CNT: resources; AT: data curation; AT: writing - original draft; AT, CS, SVA, DE, TJV, ST, and CNT: writing - review & editing; AT, and CNT: visualization; AT, and CNT: project administration; SVA, and CNT: supervision; SVA, and CNT: funding acquisition

## CONFLICT OF INTEREST

Douglas Eudy and Slobodan Trifunovic are Bayer Crop Science employees. The remaining authors declare no conflicts of interest.

## FUNDING

This study was supported by FFAR (project title: Evaluating the relative influence of maize breeding, field management, and environmental setting on crop production and sustainability targets), Bayer Crop Science, Leopold Center for Sustainable Development, the Plant Science Institute of Iowa State University, and the US. Department of Agriculture, Agricultural Research Service.

